# MitoTracker transfers from astrocytes to neurons independently of mitochondria

**DOI:** 10.1101/2025.05.27.655523

**Authors:** Katriona L. Hole, Rosalind Norkett, Emma Russell, Molly Strom, Jack H. Howden, Nicola J. Corbett, Janet Brownlees, Michael J. Devine

## Abstract

The neuroprotective transfer of mitochondria from astrocytes to neurons has been primarily investigated by labelling astrocytic mitochondria with the dye MitoTracker. Here we report that MitoTracker transfers to neurons from both astrocytes and astrocyte-conditioned media, independently of mitochondrial transfer. Our observations should prompt an essential re-evaluation of the literature concerning astrocyte-neuron mitochondrial transfer and in other systems in which contact-independent transfer has been observed using mitochondrial dyes.

## Main

Intercellular mitochondrial transfer (IMT) from astrocytes to neurons has been reported in sixteen papers from fifteen independent groups since the first report in 2016 ^1–16^.

From these reports, it has been concluded that astrocytes can release functionally intact mitochondria, either within extracellular vesicles (EV-mitochondria) or unenveloped, and neurons co-cultured with astrocytes or astrocyte-conditioned media (ACM) can take up astrocytic mitochondria. This transfer is upregulated following the application of mitochondrial stress to neurons, including oxygen/glucose deprivation ^2,10,11,15^, cisplatin ^1^, or rotenone ^9^. Furthermore, increased transfer correlates with improved neuronal viability ^1,2,9–11,15^. Notably, this neuroprotection can be observed *in vivo*, where transplantation of astrocytic mitochondria to the brains of mice can prevent neuronal damage in models of ischemia ^2,12,15^.

To study IMT, it is essential to selectively label mitochondria in the donor cell ^17,18^. For astrocyte-neuron transfer, the majority of studies have employed the mitochondrial dye MitoTracker™ (mainly CMXRos but also Green/Deep Red) to label astrocytic mitochondria ^4– 16^. However, a recent report showed that MitoTracker, which binds thiol-groups in mitochondria^19^, can transfer between macrophages, B16 cells, HEK 293T cells and immortalised bone marrow derived cells independently of mitochondrial transfer ^20^. This dye transfer was found to be dependent on cell contact in the systems that were investigated, with no transfer observed in transwell co-cultures which separate donor and acceptor cells. However, many MitoTracker-based studies have shown that astrocytic mitochondria can transfer through transwells and ACM ^4–7,11,13–15^. Therefore, it is uncertain to what extent previously reported astrocyte-neuron IMT is due to the transfer of dye rather than mitochondria. The aim of this study was to provide clarity on this issue.

To do this, we generated primary cortical astrocyte cultures from MitoTag x *GFAP-cre* mice ^21^, where outer-mitochondrial membrane-targeted GFP (GFP-OMM) is expressed exclusively in astrocytes. These astrocytes were then labelled with the mitochondrial dye MitoTracker and washed thoroughly to eliminate any residual extracellular dye (Fig. 1a, b). Using these dual mitochondrially-labelled astrocytes, we set out to compare the transfer of dye vs the transfer of mitochondria labelled with the genetically-encoded tag.

**Figure 1.**
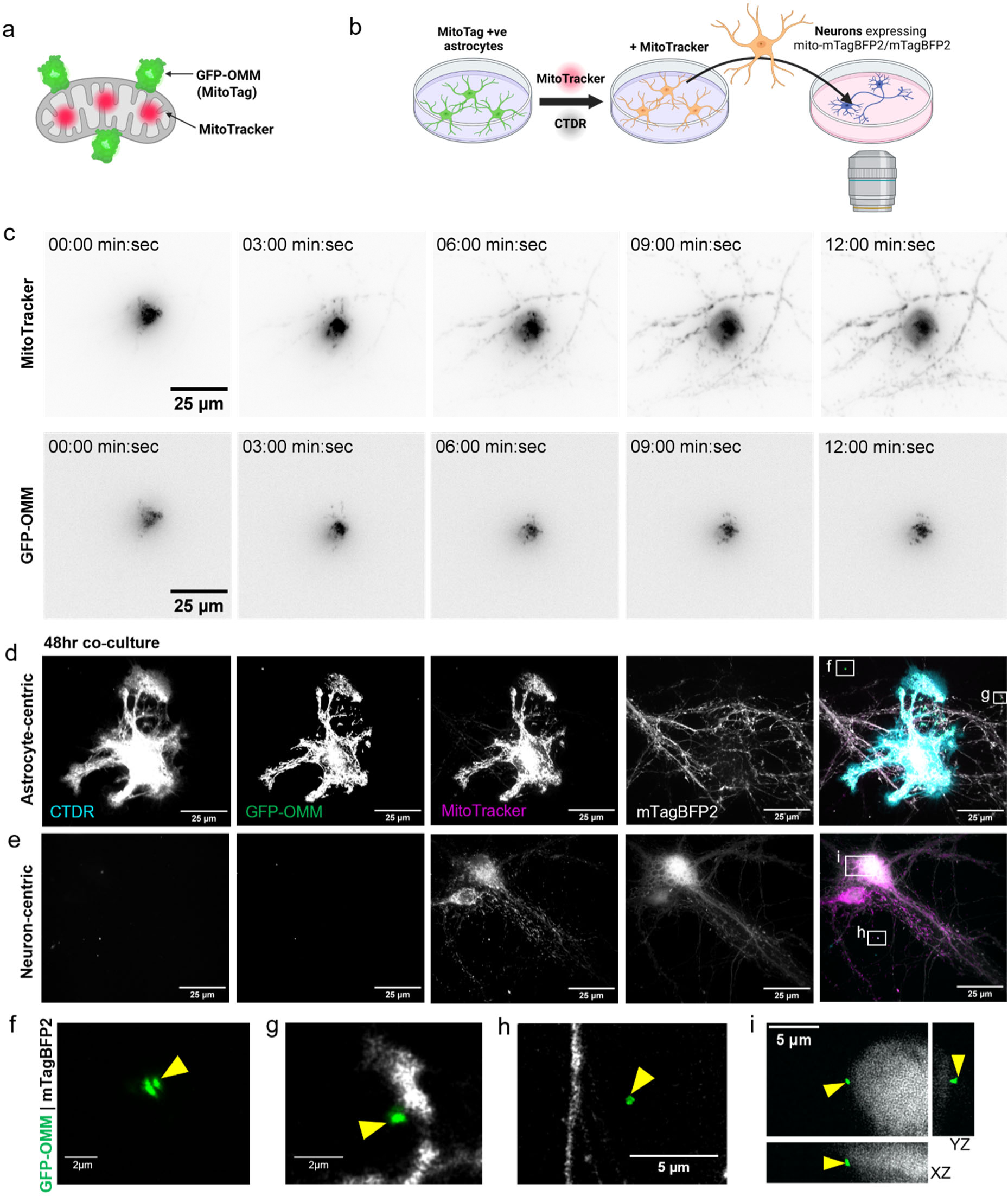
MitoTracker, but not mitochondria, transfers rapidly to neurons. (a) Mitochondria are dual labelled with outer mitochondrial membrane targeted GFP (GFP-OMM) (green) and the MitoTracker dye (red). (b) Schematic diagram outlining the protocol for live co-culture experiments. CTDR = CellTracker Deep red. (c) Timelapse images of a dual labelled astrocyte (central) immediately following co-culture with neurons. (n = 3) (d,e) Representative images after 48hr co-culture of neurons with astrocytes and immunolabelling against GFP. (f-i) Insets from (d,e) showing extracellular astrocytic mitochondria that are associated (g,i) or not associated (f,h) with neurons. All images are maximum projections except (j) which shows an orthogonal view. Yellow arrows = astrocytic mitochondria.

We first compared mitochondrial dye and MitoTag transfer in neuron-astrocyte cocultures because mitochondrial dye transfer was previously shown to be contact dependent. By imaging co-cultures immediately after application of astrocytes to primary cortical neurons expressing mTagBFP2, we observed that MitoTracker rapidly labelled mitochondria within adjacent neurons after only a few minutes (Fig. 1c, Supp. Fig. 1a, and Supp. Video 1). However, GFP-OMM, which corresponds to mitochondria, did not transfer to neurons within the acquisition period of 30 minutes. Therefore, no IMT occurred over this brief timeframe. The same observation could be made away from the site of image acquisition (Supp. Fig. 1b). After co-culture for 48 hours, a typical experimental duration based on previous reports, GFP-OMM, amplified by anti-GFP immunofluorescence, could be identified extracellularly to astrocytes (Fig. 1d-i). Most of these extracellular astrocytic mitochondria were proximal to neurons but none were transferred to neurons (Fig. 1g,i, and Supp. Fig. 1c), while the mitochondrial dye was observed within neurons (Fig. 1d,e). Therefore, we find that in contact-co-cultures mitochondrial dye transfers rapidly from astrocytes to neurons, without concomitant transfer of mitochondria.

We hypothesised that MitoTracker could also be transferred via ACM. To explore this, ACM from dual labelled astrocytes was applied to neurons during image acquisition (Fig. 2a). Notably, the MitoTracker fluorescence significantly increased in neurons throughout the image acquisition period, while the GFP-OMM signal did not increase above background (Fig. 2b-d, Supp. Video 2). Furthermore, depletion of mitochondria/EVs by filtration or further centrifugation of ACM reduced the fluorescence intensity in neurons, reflecting a reduction in mitochondrial dye transfer (Fig. 2e,f). This suggests that EVs or other cell components that are depleted by filtration or centrifugation can augment mitochondrial dye transfer ^11,17^. After 24 hours, we were still unable to detect GFP-OMM uptake within neurons whereas MitoTracker was still observable (Fig. 2g). Therefore, MitoTracker can transfer from astrocyte-conditioned media independently of mitochondrial transfer.

**Figure 2.**
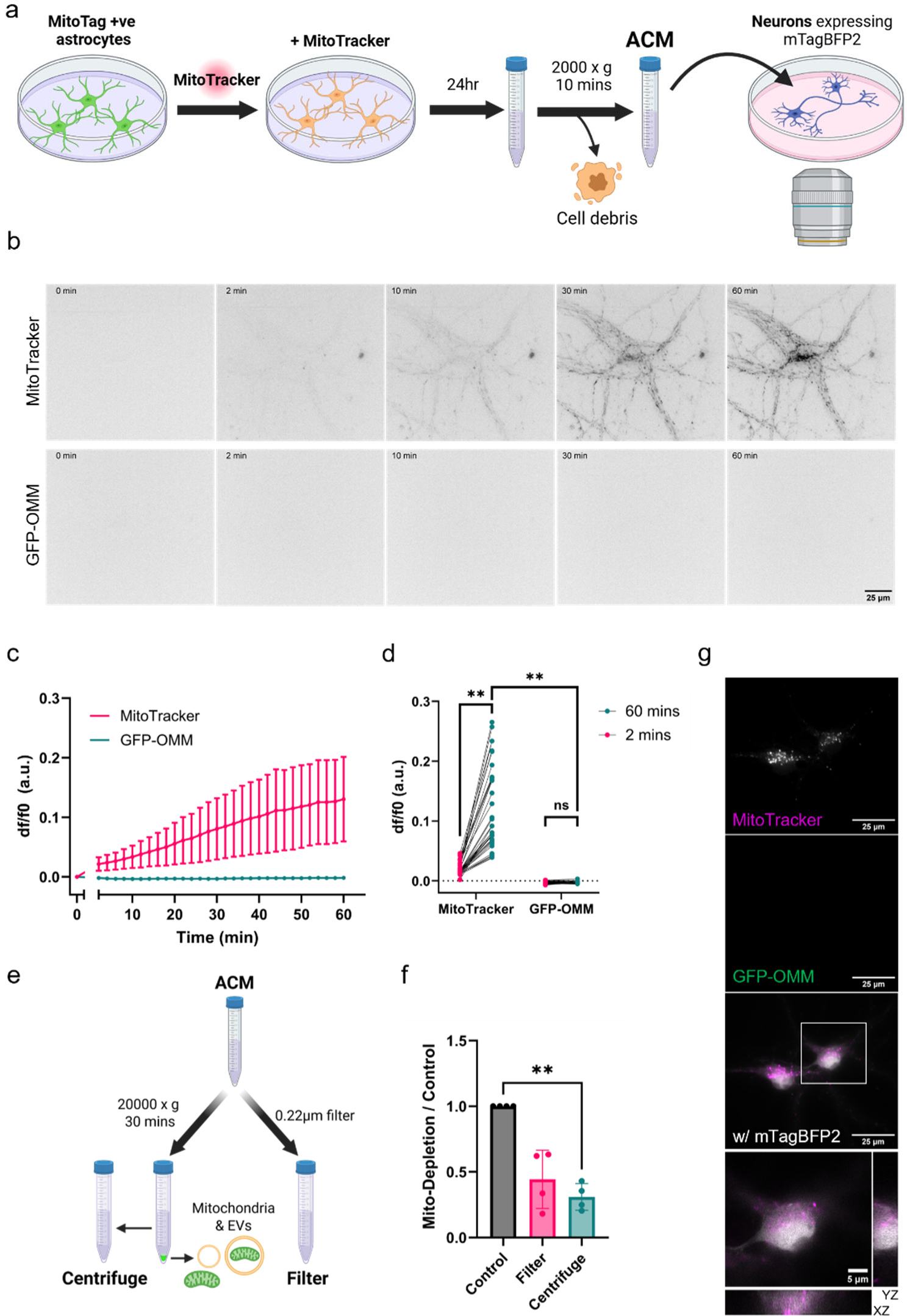
MitoTracker can transfer without direct cell-cell contact. (a) Schematic diagram outlining the protocol for live ACM incubation experiments. (b) Timelapse microscopy of neurons incubated with ACM after background has been subtracted or with increased brightness. (c) The mean somatic fluorescence intensity over time for neurons incubated with ACM, normalised to t=0. (d) Comparing the somatic fluorescence intensity at 2 mins and 60 mins for MitoTracker and GFP-OMM. Measurements from individual cells shown, statistics undertaken on biological repeats. Two-way repeated measures ANOVA with Uncorrected Fisher’s LSD, n = 4 biological repeats, 7-12 cells per repeat. (e) Schematic describing methods used to deplete mitochondria & EVs. (f) The ratio of somatic fluorescence intensity from mito-depleted ACM relative to control ACM. Ratio paired t-test, n = 4 biological repeats, 7-13 cells per repeat. (g) Representative image of neurons following 24hr incubation with astrocytes and immunolabelling against GFP with an inset showing the orthogonal view. *\*\*P* < 0.01, ns = not significant. Error bars = +/-standard deviation.

We have shown that the mitochondrial dye MitoTracker rapidly transfers from both astrocytes and ACM to neurons, independently of mitochondrial transfer. Therefore, MitoTracker transfer does not correspond to mitochondrial transfer and should not be used to investigate IMT.

We were unable to identify transfer of astrocytic mitochondria to neurons using genetically-encoded markers. However, we recognise that light microscopy has much lower throughput than flow cytometry, which is more commonly used to assess transfer. As such, we do not refute that mitochondrial transfer from astrocytes to neurons can occur. This is supported by the existing evidence for mitochondrial transfer in studies using genetically-encoded mitochondrial labels ^1–3^. Instead, we believe that IMT from astrocytes to neurons is much less common than previously reported. Furthermore, it remains possible that neuronal stress such as ischemia could increase IMT, however, this needs to be confirmed using genetically-encoded mitochondrial labels.

The release of free or EV-mitochondria by astrocytes is now well established through electron microscopy, western blots and ATP assays of conditioned media ^2,14,15^. In support of this, we were able to identify extracellular astrocytic mitochondria, some of which were in close proximity to, but not internalised by, neurons. While the work described here brings into question IMT to neurons, we do not deny the potential benefits of mitochondrial release by astrocytes. For example, there is good evidence suggesting that astrocyte conditioned media or mitochondrial transplantation can be neuroprotective, both *in vitro* and *in vivo*, against ischaemic insults ^2,11,12,15^. However, we suggest a reconsideration of whether mitochondrial transfer is itself neuroprotective, or if the presence of mitochondria and/or EVs in the extracellular space is sufficient. Alternatively, the neuroprotective effect of ACM could be elicited by EVs not containing mitochondria.

In summary, we agree that MitoTracker dye should not be used to study intercellular mitochondrial transfer ^18,20^. Moreover, it is essential that previous findings are re-examined in the context of dye transfer. Further experiments using appropriate genetically-encoded mitochondrial labels will be necessary to validate and improve the understanding of mitochondrial transfer from astrocytes to neurons.

## Acknowledgements

The authors thank the Devine laboratory for their input, particularly Jonathan Spencer for assistance with proofreading and Patrick Cottilli for assistance with astrocyte preparation. We thank Michael Way for his feedback on the manuscript. We thank the following core facilities at the Francis Crick Institute: Advanced Light Microscopy for imaging support; the Biological Research Facility for assistance with animal work; and Vector Core for plasmid and virus preparation.

This work was supported by The Francis Crick Institute, which receives its core funding from Cancer Research UK (CC2206), the UK Medical Research Council (CC2206), and the Wellcome Trust (CC2206). K.L.H was supported by funding from MSD and the MRC as part of the Crick-MSD research alliance.

Schematics and diagrams were created in BioRender. Hole, K. (2025) https://BioRender.com/olqio7s, https://BioRender.com/vas9we6, https://BioRender.com/c00spf5.

## Author Contributions

K.L.H., M.J.D, N.J.C., J.H.H., and J.B. conceived and designed the study. K.L.H. performed all the experiments and analysis. E.M. established the transgenic mouse lines. M.S. designed the plasmids and generated the viruses. R.N. assisted with experimental design and preparation of primary neurons. K.L.H. and M.J.D. wrote the original manuscript and all authors contributed to improving it.

## Methods and Materials

### Animals

Animal work was approved by the Francis Crick ethical committee and performed under UK home office licence PP3668665. All animal procedures were carried out at the Francis Crick Institute in accordance with the regulatory standards of the UK Home Office (ASPA 1986 including Amendment Regulations 2012). Mice were housed and bred under specific pathogen-free conditions (SPF) in individually ventilated cages under a 12hr light–dark cycle at ambient temperature (19°C-21°C) and humidity (45-55%). Standard food and water were provided *ad libitum*. Additional information can be found in Supplementary Information Table 1. Neurons were derived from E16.5 embryos of C57BL/6J RRID:IMSR_JAX:000664, mice of either sex. B6.Cg-Tg(Gfap-cre)77.6Mvs/2J - RRID:IMSR_JAX:024098 (GFAP-Cre) mice and B6N.Cg-Gt(ROSA)26Sortm1(CAG-EGFP*)Thm/J - RRID:IMSR_JAX:032675 (MitoTag) mice were separately rederived to C57BL/6J mice and backcrossed for 2 and 8 generations respectively, before crossing to generate *GFAP*-cre x MitoTag mice. The breeding strategy was optimised to account for known potential occurrence of germline deletion of the floxed allele when breeding from males as previously reported ^22^ – only female *GFAP*-cre +ve mice were used for breeding. Astrocytes were generated from P0-2 MitoTag x *GFAP*-cre pups of either sex.

### Cell Culture

#### Primary neuron culture

Primary cortical neurons were prepared as previously described ^23^. All materials are from Gibco unless stated otherwise. The day before culture, glass coverslips (GG-25-1.5H-Pre, Neuvitro) were pre-coated overnight with 0.5mg/mL PLL at 37°C. The following day, coverslips were washed twice in dH2O and left in attachment media (10% heat inactivated horse serum, 1mM sodium pyruvate, 33mM Glucose in MEM) at 37°C, 5% CO2 until use.

E16.5 embryos were harvested and tissue was kept in ice-cold HBSS (1X HBSS, 10mM HEPES pH7.3, in H_2_O) throughout the dissection. Following decapitation, the brain was removed from the skull and the hemispheres separated. The cortices were dissected and then incubated in 0.05% trypsin containing 10μg/mL DNase I (Sigma) for 15 minutes at 37°C, 5% CO2 with gentle agitation every 2-3 minutes. The trypsin was removed and the cortices washed three times in HBSS. Tissue was then triturated 15 times in attachment media containing 10μg/mL DNase I (Sigma) with a P1000 pipette. Dissociated cells were transferred to a new 15mL falcon tube. Cells were counted before plating on the pre-coated coverslips at a density of 125,000 cells per well. 5 hours post-plating, media was replaced with maintenance media (2% B27, 1% GlutaMAX, 33mM glucose in Neurobasal). From DIV5, a half media change was undertaken with BrainPhys™-Neurocult™ SM1 (STEMCELL Technologies) every 2-3 days until use at DIV12-14.

#### Adeno-associated virus infection

AAV-hSyn-mTagBFP2 and AAV-hSyn-mito-mTagBFP2, with AAV2/1 serotype, were created by The Francis Crick Vector Core facility at the Francis Crick Institute. Sequences can be found in supplementary information. pAAV2/1 was a gift from James M. Wilson (Addgene plasmid #112862; http://n2t.net/addgene:112862; RRID:Addgene_112862) Neurons were infected at DIV6/7 and allowed to express for 6-7 days prior to experiments.

#### Primary astrocyte culture

Primary cortical astrocytes were derived from P0-2 homozygous MitoTag x *GFAP*-cre mice of either sex and isolated as previously described ^24^. Tail tissue was taken for genotyping by Transnetyx. Cortices were incubated in papain solution (20 U/mL papain (Worthington Biochemicals, cat. LS003124) in Hibernate-A) for 15 minutes at 37°C, 5% CO2, then washed three times PBS-Glucose (0.585% glucose in PBS). The tissue was triturated with an FCS-coated P1000 pipette in astrocyte media (DMEM, cat. 41966-029 with 10% FCS) with 10μg/mL DNase I (Sigma). Astrocytes were then centrifuged at 750 x g for 5 mins and the pellet resuspended in astrocyte media. Cells were plated in uncoated T75 flasks at 1 × 10^6^ cells and incubated at 37°C, 5% CO2. The media was replaced after 24 hours and every 2-3 days after until the cells reached confluency. At this point, astrocytes were trypsinised, centrifuged at 300 x g for 10 minutes and resuspended in recovery cell culture freezing medium (Gibco, 11560446) at 4M cells per mL. 0.5mL astrocytes were transferred to cryovials and stored in a Mr. Frosty™ freezing container at −70°C for at least 24 hours then transferred to liquid nitrogen for long term storage. When needed, astrocytes were thawed and plated in astrocyte media at 300,000 cells in 60mm dishes. These cells were maintained as before until use.

#### Primary astrocyte-neuron co-cultures

Astrocytes for co-culture were labelled with 100nM MitoTracker™ Red CMXRos (Invitrogen, cat. M7512) and CellTracker Deep Red (1:1000 dilution, CTDR, Invitrogen, cat. C34565) in unsupplemented DMEM for 30 mins. Following a single wash in astrocyte media, astrocytes were washed five times with PBS to ensure removal of extracellular dye. Astrocytes were then immediately trypsinized and the cell suspension centrifuged at 300 x g for 10 mins before resuspension in BrainPhys-SM1 media. Astrocytes were added to neurons at a ratio of 1:1, keeping neuronal conditioned media at 50%.

For immunofluorescence experiments, co-cultures were fixed after 48-hours. For live imaging experiments, astrocytes were added to neurons in the imaging chamber, positions of interest identified and images acquired within 3 mins of application.

#### Astrocyte conditioned media

24hr prior to imaging, astrocytes were labelled with MitoTracker™ CMXRos as described above, including the washing steps. Astrocytes were then cultured in BrainPhys-SM1 for 24 hours before the astrocyte conditioned media was collected and centrifuged at 2000 x g for 10 mins to eliminate any cells or debris present in the media, with the supernatant kept as ACM. For mitochondrial depletion, the ACM was either passed through a 0.22μm filter or centrifuged at 20,000 x g for 30 mins to pellet mitochondria and extracellular vesicles, retaining the supernatant.

For fixed imaging experiments, 1mL neuronal media was replaced with 1mL ACM, and neurons were incubated for 24 hours prior to fixing.

For live imaging experiments, neurons were initially imaged in 500μL neuronal conditioned media (t = 0), and then 500μL ACM was applied during acquisition. Images were acquired every 2 mins for 62 mins, with z-stacks of 21 × 0.5μm steps.

#### Immunocytochemistry

Fixing solution (4% paraformaldehyde (Electron Microscopy Sciences, 1570), 4% sucrose in PBS) was prewarmed to 37°C before incubating with cells for 15 mins. Following 3 × 5 min washes with PBS, cells were left in PBS at 4°C before use. Cells were permeabilised with 0.1% Triton-X-100 in PBS for 5 mins, followed by 3 x PBS washes. Cells were blocked for 30 minutes in blocking buffer (1% BSA (Sigma), 1% horse serum (Gibco) in PBS). Antibodies were diluted in blocking buffer as follows: anti-ATPB (abcam, ab14730, 1:500), anti-GFP (abcam, ab6556, 1:2000). Coverslips were incubated with primary antibodies for 30 mins, washed 3 × 5 mins with blocking buffer, then incubated with secondary antibodies for 30 mins: Chicken anti-Rabbit IgG (H+L) Cross-Adsorbed Secondary Antibody, Alexa Fluor™ 488 (Invitrogen, A-21441, 1:1000); Donkey anti-Mouse IgG (H+L) Highly Cross-Adsorbed Secondary Antibody, Alexa Fluor™ 647 (Invitrogen, A-31571, 1:1000). Coverslips were washed for a further 3 × 5 mins in blocking buffer before a final wash step with for 3 × 5 mins in PBS. Coverslips were mounted onto slides using ProLong Glass Antifade Mountant and allowed to cure for 24 hours at rtp before sealing with nail varnish.

### Microscopy and image analysis

Confocal imaging was undertaken using a Visitech iSIM microscope on an IX83 microscope body and widefield microscopy and Micro-Manager v2.1.0 software ^25^. For live imaging, a 60X, 1.4 NA objective was used. Post-timelapse and fixed imaging was undertaken with a 100X, 1.5 NA objective. Live imaging was undertaken at 37°C with 5% CO_2_.

Widefield microscopy was undertaken with a fluorescent microscope on a Nikon Ti2 microscope body (Evident) with a 20X, 0.75NA objective.

All image analysis was undertaken using Fiji (2.16.0) ^26^. Images from fixed samples were deconvoluted using Microvolution® software. To measure fluorescence intensity, a sum-stack z-projection was created. The cell body was outlined using the cell-fill as reference. The mean fluorescence intensity within that area was measured over time for each channel of interest. For df/f0 calculations, f0 relates to the fluorescence before ACM was added. Where necessary, drift correction was undertaken using the Correct 3D Drift plug-in ^27^. Statistics was calculated using GraphPad Prism 10.4.1.

