## Supplementary Information for "MitoTracker transfers from astrocytes to neurons independently of mitochondria"

**Supplementary Video 1. MitoTracker labels adjacent neuronal mitochondria without mitochondrial transfer.** Image acquisition immediately after astrocytes expressing GFP-OMM (green) and labelled with MitoTracker dye (magenta) are co-cultured with neurons expressing mito-mTagBFP2 (blue).

**Supplementary Video 2. MitoTracker transfers from ACM to neurons without mitochondria.** Neurons expressing mTagBFP2 (blue) are incubated with ACM from astrocytes expressing GFP-OMM (green) and labelled with MitoTracker dye (magenta).

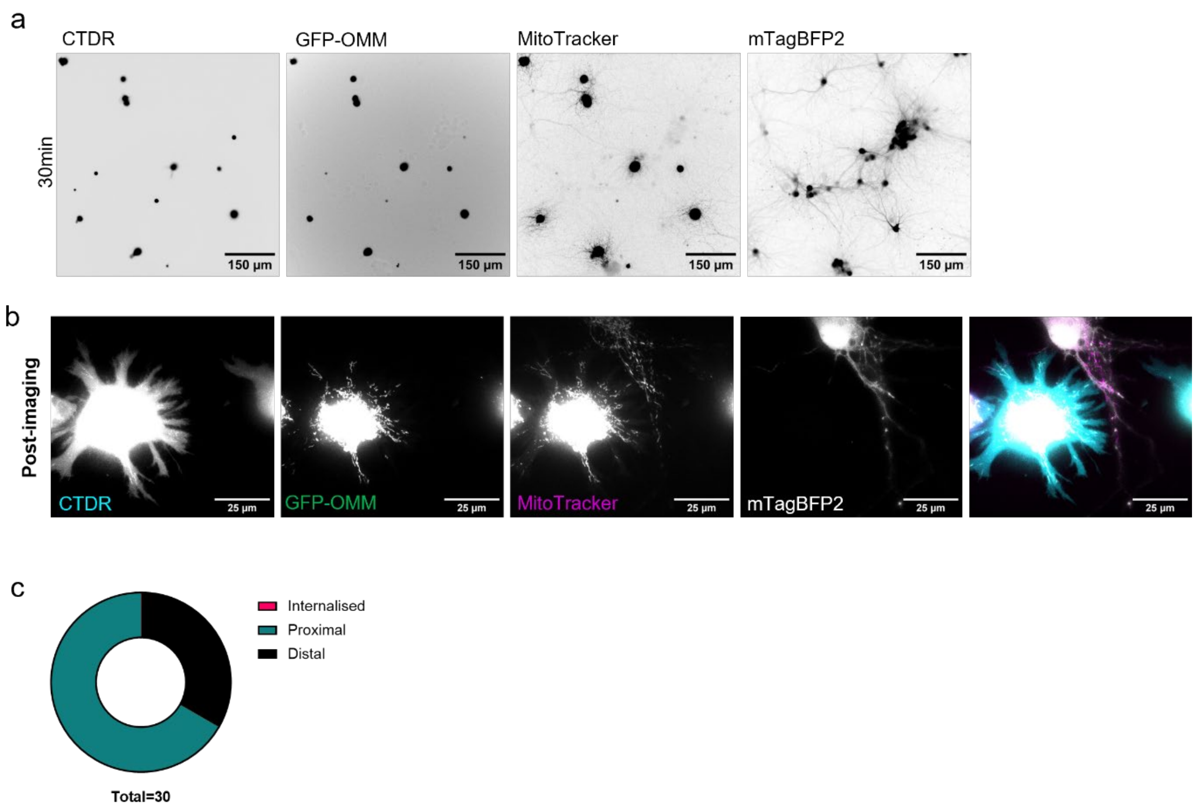


**Extended Data 1. Contact-dependent transfer of MitoTracker but not mitochondria** (a) Live imaging of mTagBFP2 expressing neurons co-cultured with dual-mitochondrially labelled astrocytes for 30 minutes imaged by widefield microscopy. MitoTracker transferred to adjacent neurons first. (b) Representative images of neuron-astrocyte co-cultures immediately following timelapse imaging in a different, unimaged, region (n = 3). Of the 30 extracellular-astrocytic mitochondria identified in one imaging session of 48hr co-cultures, 20 were proximal to neurons, 10 were not associated with neurons and none were internalised by neurons.

Supplementary Information Table 1: Details of housing and husbandry for mice.

| Categories | Details |
|---|---|
| **Cage/tank/housing system (type and** | Individually Ventilated Cages Green line, floor area 500 cm2 (Tecniplast, Italy), Isolators |
| **Food and water (type, composition, supplier and access)** | 2018 Teklad global diet (Envigo, UK) autoclaved before use;<br>Isolators: T.2918CSD irradiated diet (Envigo, UK)<br>Drinking water is mains water passed through RO system and provided in cages by Automated Watering System (Avidity Science, USA) |
| **Bedding and nesting material** | Bedding: Aspen 4HK (Datesand, UK)<br>Nesting material: Bed R nest (Datesand, UK) |
| **Temperature and humidity** | Temperature 21oC ± 2oC (in May 2022 temperature in holding rooms was raised to 22 +/- 2)<br>Humidity: 55% RH ± 10% |
| **Sanitation (frequency of cage/tank water changes, material transferred, water quality)** | Cages are changed as required following SOP on discretionary cage changing with some enrichment and nesting material transferred to new cage to minimise animal stress.<br>Cage cleaning is done via a Tecniplast Pegasus robotic & tunnel washing system and then autoclaved before use. Cage lids and food hoppers are processed via rack washer and then autoclaved as and when required.<br>Avidity automatic watering system is flushed once per day. Animal drinking water is RO treated water with addition of chlorine. |
| **Social environment (group size and composition/stocking density)** | Mice are group housed with maximum occupancy in accordance with regulatory requirements. Mice with body weight >20g are housed with up to 5 mice per cage. Breeding set ups are pairs and trios. Experimental breeding pairs (to produce pups for primary astrocytes) were separated and the mother housed singly once plugged. Mice are only individually housed singly if required by experiment, for welfare reasons, or if no alternative was available. |
| **Biosecurity (level)** | A hybrid health monitoring programme which utilizes dirty bedding sentinels and environmental swabs is employed.<br>For mice, at any one time two sentinels are in place per group of racks and are taken for examination when they have been exposed to the environment for approximately six months. Environmental swabs are taken from surfaces exposed to exhaust air from the racks every 4 months.<br>Blood is drawn from sentinels, once only, for serology during the monitoring period. At the end of the exposure period the sentinels are killed by an i.p. overdose of pentobarbitone and subjected to a full necropsy. The pelt is examined for ectoparasites and samples taken include blood, throat and caecal swabs, and faecal pellets. Other swabs and tissues are taken are when deemed necessary. Wet mounts of gut contents are examined for endoparasites and tape tests performed for Syphacia. Gross abnormalities and lesions are recorded and investigated further as necessary.<br>Bacteriology, serology and basic parasitology is carried out by the Crick BRF microbiology laboratory. Some other analyses are outsourced. The choice of agents screened and screening frequencies conform broadly to FELASA recommendations, with some amendments as deemed appropriate by the Veterinary and Animal Health Services Team |
| **Lighting (type, schedule and intensity)** | 12 hrs day /night cycles with 70% light intensity with 15 minutes gradual increase/ decrease in light intensity |
| **Environmental enrichment** | Cage balconies (Tecniplast, UK), and cardboard mouse houses (Datesand, UK) provided. |
| **Sex of the animals** | Pups and embryos of either sex were used. |

**Plasmid sequences**

pAAV hSyn mito-mTagBFP2

ITRs=green

hSynapsin promoter =Red

COX8=pink

mTagBFP2=blue

WPRE=orange

hGH=purple

acatgtcctgcaggcagctgcgcgctcgctcgctcactgaggccgcccgggcgtcgggcgacctttggtcgcccggcctcagtgagcg
agcgagcgcgcagagagggagtggccaactccatcactaggggttcctgcggccgcacgcgtgtgtctagactgcagagggccctg
cgtatgagtgcaagtgggttttaggaccaggatgaggcggggtgggggtgcctacctgacgaccgaccccgacccactggacaagc
acccaacccccattccccaaattgcgcatcccctatcagagagggggaggggaaacaggatgcggcgaggcgcgtgcgcactgcca
gcttcagcaccgcggacagtgccttcgcccccgcctggcggcgcgcgccaccgccgcctcagcactgaaggcgcgctgacgtcactc
gccggtcccccgcaaactccccttcccggccaccttggtcgcgtccgcgccgccgccggcccagccggaccgcaccacgcgaggcgc
gagataggggggcacgggcgcgaccatctgcgctgcggcgccggcgactcagcgctgcctcagtctgcggtgggcagcggaggag
tcgtgtcgtgcctgagagcgcagtcgagaaggtaccggatcctctagagtcgacgccaccatgtccgtcctgacgccgctgctgctgc
ggggcttgacaggctcggcccggcggctcccagtgccgcgcgccaagatccattcgttgggggatccaccggtatgagcgagctgat
taaggagaacatgcacatgaagctgtacatggagggcaccgtggacaaccatcacttcaagtgcacatccgagggcgaaggcaag
ccctacgagggcacccagaccatgagaatcaaggtggtcgagggcggccctctccccttcgccttcgacatcctggctactagcttcct
ctacggcagcaagaccttcatcaaccacacccagggcatccccgacttcttcaagcagtccttccctgagggcttcacatgggagaga
gtcaccacatacgaagacgggggcgtgctgaccgctacccaggacaccagcctccaggacggctgcctcatctacaacgtcaagat
cagaggggtgaacttcacatccaacggccctgtgatgcagaagaaaacactcggctgggaggccttcaccgagacgctgtaccccg
ctgacggcggcctggaaggcagaaacgacatggccctgaagctcgtgggcgggagccatctgatcgcaaacgccaagaccacata
tagatccaagaaacccgctaagaacctcaagatgcctggcgtctactatgtggactacagactggaaagaatcaaggaggccaaca
acgagacctacgtcgagcagcacgaggtggcagtggccagatactgcgacctccctagcaaactggggcacaagcttaattaagaa
ttcgatatcaagcttatcgataatcaacctctggattacaaaatttgtgaaagattgactggtattcttaactatgttgctccttttacgcta
tgtggatacgctgctttaatgcctttgtatcatgctattgcttcccgtatggctttcattttctcctccttgtataaatcctggttgctgtctctt
tatgaggagttgtggcccgttgtcaggcaacgtggcgtggtgtgcactgtgtttgctgacgcaacccccactggttggggcattgccac
cacctgtcagctcctttccgggactttcgctttccccctccctattgccacggcggaactcatcgccgcctgccttgcccgctgctggaca
ggggctcggctgttgggcactgacaattccgtggtgttgtcggggaaatcatcgtcctttccttggctgctcgcctatgttgccacctgg
attctgcgcgggacgtccttctgctacgtcccttcggccctcaatccagcggaccttccttcccgcggcctgctgccggctctgcggcctc
ttccgcgtcttcgccttcgccctcagacgagtcggatctccctttgggccgcctccccgcatcgataccgagcgctgctcgagagatcta
cgggtggcatccctgtgacccctccccagtgcctctcctggccctggaagttgccactccagtgcccaccagccttgtcctaataaaatt
aagttgcatcattttgtctgactaggtgtccttctataatattatggggtggaggggggtggtatggagcaaggggcaagttgggaaga
caacctgtagggcctgcggggtctattgggaaccaagctggagtgcagtggcacaatcttggctcactgcaatctccgcctcctgggt
tcaagcgattctcctgcctcagcctcccgagttgttgggattccaggcatgcatgaccaggctcagctaattttgtttttttggtagagac
ggggtttcaccatattggccaggctggtctccaactcctaatctcaggtgatctacccaccttggcctcccaaattgctgggattacagg
cgtgaaccactgctcccttccctgtccttctgattttgtaggtaaccacgtgcggaccgagcggccgcaggaacccctagtgatggagt
tggccactccctctctgcgcgctcgctcgctcactgaggccgggcgaccaaaggtcgcccgacgcccgggctttgcccgggcggcctc
agtgagcgagcgagcgcgcagctgcctgcaggggcgcctgatgcggtattttctccttacgcatctgtgcggtatttcacaccgcatac
gtcaaagcaaccatagtacgcgccctgtagcggcgcattaagcgcggcgggtgtggtggttacgcgcagcgtgaccgctacacttgc
cagcgccttagcgcccgctcctttcgctttcttcccttcctttctcgccacgttcgccggctttccccgtcaagctctaaatcgggggctcc
ctttagggttccgatttagtgctttacggcacctcgaccccaaaaaacttgatttgggtgatggttcacgtagtgggccatcgccctgat

agacggtttttcgccctttgacgttggagtccacgttctttaatagtggactcttgttccaaactggaacaacactcaactctatctcggg
ctattcttttgatttataagggattttgccgatttcggtctattggttaaaaaatgagctgatttaacaaaaatttaacgcgaattttaaca
aaatattaacgtttacaattttatggtgcactctcagtacaatctgctctgatgccgcatagttaagccagccccgacacccgccaacac
ccgctgacgcgccctgacgggcttgtctgctcccggcatccgcttacagacaagctgtgaccgtctccgggagctgcatgtgtcagag
gttttcaccgtcatcaccgaaacgcgcgagacgaaagggcctcgtgatacgcctatttttataggttaatgtcatgataataatggtttct
tagacgtcaggtggcacttttcggggaaatgtgcgcggaacccctatttgtttatttttctaaatacattcaaatatgtatccgctcatga
gacaataaccctgataaatgcttcaataatattgaaaaaggaagagtatgagtattcaacatttccgtgtcgcccttattcccttttttgc
ggcattttgccttcctgtttttgctcacccagaaacgctggtgaaagtaaaagatgctgaagatcagttgggtgcacgagtgggttacat
cgaactggatctcaacagcggtaagatccttgagagttttcgccccgaagaacgttttccaatgatgagcacttttaaagttctgctatg
tggcgcggtattatcccgtattgacgccgggcaagagcaactcggtcgccgcatacactattctcagaatgacttggttgagtactcac
cagtcacagaaaagcatcttacggatggcatgacagtaagagaattatgcagtgctgccataaccatgagtgataacactgcggcca
acttacttctgacaacgatcggaggaccgaaggagctaaccgcttttttgcacaacatgggggatcatgtaactcgccttgatcgttgg
gaaccggagctgaatgaagccataccaaacgacgagcgtgacaccacgatgcctgtagcaatggcaacaacgttgcgcaaactatt
aactggcgaactacttactctagcttcccggcaacaattaatagactggatggaggcggataaagttgcaggaccacttctgcgctcg
gcccttccggctggctggtttattgctgataaatctggagccggtgagcgtgggtctcgcggtatcattgcagcactggggccagatgg
taagccctcccgtatcgtagttatctacacgacggggagtcaggcaactatggatgaacgaaatagacagatcgctgagataggtgc
ctcactgattaagcattggtaactgtcagaccaagtttactcatatatactttagattgatttaaaacttcatttttaatttaaaaggatcta
ggtgaagatcctttttgataatctcatgaccaaaatcccttaacgtgagttttcgttccactgagcgtcagaccccgtagaaaagatcaa
aggatcttcttgagatcctttttttctgcgcgtaatctgctgcttgcaaacaaaaaaaccaccgctaccagcggtggtttgtttgccggat
caagagctaccaactctttttccgaaggtaactggcttcagcagagcgcagataccaaatactgttcttctagtgtagccgtagttaggc
caccacttcaagaactctgtagcaccgcctacatacctcgctctgctaatcctgttaccagtggctgctgccagtggcgataagtcgtgt
cttaccgggttggactcaagacgatagttaccggataaggcgcagcggtcgggctgaacggggggttcgtgcacacagcccagcttg
gagcgaacgacctacaccgaactgagatacctacagcgtgagctatgagaaagcgccacgcttcccgaagggagaaaggcggaca
ggtatccggtaagcggcagggtcggaacaggagagcgcacgagggagcttccagggggaaacgcctggtatctttatagtcctgtc
gggtttcgccacctctgacttgagcgtcgatttttgtgatgctcgtcaggggggcggagcctatggaaaaacgccagcaacgcggcctt
tttacggttcctggccttttgctggccttttgctc

pAAV hSyn mTagBFP2

ITRs=green

hSynapsin promoter =Red

mTagBFP2=blue

WPRE=orange

hGH=purple

acatgtcctgcaggcagctgcgcgctcgctcgctcactgaggccgcccgggcgtcgggcgacctttggtcgcccggcctcagtgagcg
agcgagcgcgcagagagggagtggccaactccatcactaggggttcctgcggccgcacgcgtgtgtctagactgcagagggccctg
cgtatgagtgcaagtgggttttaggaccaggatgaggcggggtgggggtgcctacctgacgaccgaccccgacccactggacaagc
acccaacccccattccccaaattgcgcatcccctatcagagagggggaggggaaacaggatgcggcgaggcgcgtgcgcactgcca
gcttcagcaccgcggacagtgccttcgcccccgcctggcggcgcgcgccaccgccgcctcagcactgaaggcgcgctgacgtcactc
gccggtcccccgcaaactccccttcccggccaccttggtcgcgtccgcgccgccgccggcccagccggaccgcaccacgcgaggcgc
gagataggggggcacgggcgcgaccatctgcgctgcggcgccggcgactcagcgctgcctcagtctgcggtgggcagcggaggag
tcgtgtcgtgcctgagagcgcagtcgagaaggtaccgccaccatgagcgagctgattaaggagaacatgcacatgaagctgtacatg
gagggcaccgtggacaaccatcacttcaagtgcacatccgagggcgaaggcaagccctacgagggcacccagaccatgagaatca
aggtggtcgagggcggccctctccccttcgccttcgacatcctggctactagcttcctctacggcagcaagaccttcatcaaccacacc
cagggcatccccgacttcttcaagcagtccttccctgagggcttcacatgggagagagtcaccacatacgaagacgggggcgtgctg
accgctacccaggacaccagcctccaggacggctgcctcatctacaacgtcaagatcagaggggtgaacttcacatccaacggccct
gtgatgcagaagaaaacactcggctgggaggccttcaccgagacgctgtaccccgctgacggcggcctggaaggcagaaacgaca
tggccctgaagctcgtgggcgggagccatctgatcgcaaacgccaagaccacatatagatccaagaaacccgctaagaacctcaag
atgcctggcgtctactatgtggactacagactggaaagaatcaaggaggccaacaacgagacctacgtcgagcagcacgaggtggc
agtggccagatactgcgacctccctagcaaactggggcacaagcttaattaagaattcgatatcaagcttatcgataatcaacctctg
gattacaaaatttgtgaaagattgactggtattcttaactatgttgctccttttacgctatgtggatacgctgctttaatgcctttgtatcat
gctattgcttcccgtatggctttcattttctcctccttgtataaatcctggttgctgtctctttatgaggagttgtggcccgttgtcaggcaac
gtggcgtggtgtgcactgtgtttgctgacgcaacccccactggttggggcattgccaccacctgtcagctcctttccgggactttcgcttt
ccccctccctattgccacggcggaactcatcgccgcctgccttgcccgctgctggacaggggctcggctgttgggcactgacaattccg
tggtgttgtcggggaaatcatcgtcctttccttggctgctcgcctatgttgccacctggattctgcgcgggacgtccttctgctacgtccct
tcggccctcaatccagcggaccttccttcccgcggcctgctgccggctctgcggcctcttccgcgtcttcgccttcgccctcagacgagtc
ggatctccctttgggccgcctccccgcatcgataccgagcgctgctcgagagatctacgggtggcatccctgtgacccctccccagtgc
ctctcctggccctggaagttgccactccagtgcccaccagccttgtcctaataaaattaagttgcatcattttgtctgactaggtgtccttc
tataatattatggggtggaggggggtggtatggagcaaggggcaagttgggaagacaacctgtagggcctgcggggtctattggga
accaagctggagtgcagtggcacaatcttggctcactgcaatctccgcctcctgggttcaagcgattctcctgcctcagcctcccgagtt
gttgggattccaggcatgcatgaccaggctcagctaattttgtttttttggtagagacggggtttcaccatattggccaggctggtctcc
aactcctaatctcaggtgatctacccaccttggcctcccaaattgctgggattacaggcgtgaaccactgctcccttccctgtccttctga
ttttgtaggtaaccacgtgcggaccgagcggccgcaggaacccctagtgatggagttggccactccctctctgcgcgctcgctcgctca
ctgaggccgggcgaccaaaggtcgcccgacgcccgggctttgcccgggcggcctcagtgagcgagcgagcgcgcagctgcctgca
ggggcgcctgatgcggtattttctccttacgcatctgtgcggtatttcacaccgcatacgtcaaagcaaccatagtacgcgccctgtagc
ggcgcattaagcgcggcgggtgtggtggttacgcgcagcgtgaccgctacacttgccagcgccttagcgcccgctcctttcgctttcttc
ccttcctttctcgccacgttcgccggctttccccgtcaagctctaaatcgggggctccctttagggttccgatttagtgctttacggcacct
cgaccccaaaaaacttgatttgggtgatggttcacgtagtgggccatcgccctgatagacggtttttcgccctttgacgttggagtccac
gttctttaatagtggactcttgttccaaactggaacaacactcaactctatctcgggctattcttttgatttataagggattttgccgatttc
ggtctattggttaaaaaatgagctgatttaacaaaaatttaacgcgaattttaacaaaatattaacgtttacaatttatggtgcactctc
agtacaatctgctctgatgccgcatagttaagccagccccgacacccgccaacacccgctgacgcgccctgacgggcttgtctgctcc
cggcatccgcttacagacaagctgtgaccgtctccgggagctgcatgtgtcagaggttttcaccgtcatcaccgaaacgcgcgagacg
aaagggcctcgtgatacgcctatttttataggttaatgtcatgataataatggtttcttagacgtcaggtggcacttttcggggaaatgtg

cgcggaacccctatttgtttatttttctaaatacattcaaatatgtatccgctcatgagacaataaccctgataaatgcttcaataatattg
aaaaaggaagagtatgagtattcaacatttccgtgtcgcccttattcccttttttgcggcattttgccttcctgtttttgctcacccagaaac
gctggtgaaagtaaaagatgctgaagatcagttgggtgcacgagtgggttacatcgaactggatctcaacagcggtaagatccttga
gagttttcgccccgaagaacgttttccaatgatgagcacttttaaagttctgctatgtggcgcggtattatcccgtattgacgccgggcaa
gagcaactcggtcgccgcatacactattctcagaatgacttggttgagtactcaccagtcacagaaaagcatcttacggatggcatga
cagtaagagaattatgcagtgctgccataaccatgagtgataacactgcggccaacttacttctgacaacgatcggaggaccgaagg
agctaaccgcttttttgcacaacatgggggatcatgtaactcgccttgatcgttgggaaccggagctgaatgaagccataccaaacga
cgagcgtgacaccacgatgcctgtagcaatggcaacaacgttgcgcaaactattaactggcgaactacttactctagcttcccggcaa
caattaatagactggatggaggcggataaagttgcaggaccacttctgcgctcggcccttccggctggctggtttattgctgataaatc
tggagccggtgagcgtgggtctcgcggtatcattgcagcactggggccagatggtaagccctcccgtatcgtagttatctacacgacg
gggagtcaggcaactatggatgaacgaaatagacagatcgctgagataggtgcctcactgattaagcattggtaactgtcagaccaa
gtttactcatatatactttagattgatttaaaacttcatttttaatttaaaaggatctaggtgaagatcctttttgataatctcatgaccaaa
atcccttaacgtgagttttcgttccactgagcgtcagaccccgtagaaaagatcaaaggatcttcttgagatcctttttttctgcgcgtaat
ctgctgcttgcaaacaaaaaaaccaccgctaccagcggtggtttgtttgccggatcaagagctaccaactctttttccgaaggtaactg
gcttcagcagagcgcagataccaaatactgttcttctagtgtagccgtagttaggccaccacttcaagaactctgtagcaccgcctaca
tacctcgctctgctaatcctgttaccagtggctgctgccagtggcgataagtcgtgtcttaccgggttggactcaagacgatagttaccg
gataaggcgcagcggtcgggctgaacggggggttcgtgcacacagcccagcttggagcgaacgacctacaccgaactgagatacct
acagcgtgagctatgagaaagcgccacgcttcccgaagggagaaaggcggacaggtatccggtaagcggcagggtcggaacagg
agagcgcacgagggagcttccagggggaaacgcctggtatctttatagtcctgtcgggtttcgccacctctgacttgagcgtcgatttt
tgtgatgctcgtcaggggggcggagcctatggaaaaacgccagcaacgcggcctttttacggttcctggccttttgctggccttttgctc
